# Human HspB1, HspB3, HspB5 and HspB8: Shaping these Disease Factors during Vertebrate Evolution

**DOI:** 10.1101/2022.02.24.481792

**Authors:** Rainer Benndorf, Ryan Velazquez, Jordan D. Zehr, Sergei Kosakovsky Pond, Jody L. Martin, Alexander G. Lucaci

## Abstract

Small heat shock proteins (sHSPs) emerged early in evolution and occur in all domains of life and nearly in all species, including humans. Mutations in four sHSPs (HspB1, HspB3, HspB5, HspB8) are associated with neuromuscular disorders. The aim of this study is to investigate the evolutionary forces shaping these sHSPs during vertebrate evolution. We performed comparative evolutionary analyses on a set of orthologous sHSP sequences, based on the ratio of non-synonymous: synonymous substitution rates for each codon. We found that these sHSPs had been historically exposed to different degrees of purifying selection, decreasing in this order: HspB8 > HspB1, HspB5 > HspB3. Within each sHSP, regions with different degrees of purifying selection can be discerned, resulting in characteristic selective pressure profiles. The conserved α-crystallin domains were exposed to the most stringent purifying selection compared to the flanking regions, supporting a ‘dimorphic pattern’ of evolution. Thus, during vertebrate evolution the different sequence partitions were exposed to different and measurable degrees of selective pressures. Among the disease-associated mutations, most are missense mutations primarily in HspB1 and to a minor extent in the other sHSPs. Our data provide an explanation for this disparate incidence. Contrary to the expectation, most missense mutations cause dominant disease phenotypes. Theoretical considerations support a connection between the historic exposure of these sHSP genes to a high degree of purifying selection and the unusual prevalence of genetic dominance of the associated disease phenotypes. Our study puts the genetics of inheritable sHSP-borne diseases into the context of vertebrate evolution.

## Introduction

The human genome encodes ten small heat shock proteins (sHSP) (Fontaine et al. 2003; Kappé et al. 2003), now systematically named HspB1 through HspB10, whether or not their expression is induced by stress factors (Kampinga et al. 2009). The defining feature of sHSPs is a conserved ~85 amino acid long sequence stretch called the α-crystallin domain (αCD) (de Jong et al. 1998; Kappé et al. 2010; Mymrikov et al. 2011). It is believed that the primary function of sHSPs is to act as holdase chaperones with a depot function for unfolded proteins, thereby shaping protein-protein interactions. This, in turn, controls many cellular processes and structures such as cell signaling, cell architecture, differentiation, redox homeostasis, apoptosis, and others (Carra et al. 2019).

sHSPs occur in all domains of life, including *Archea*, *Bacteria*, and *Eucarya*, suggesting that they emerged early in evolution. Nearly all organisms contain genes encoding sHSPs except a few pathogens that have lost these genes (Kriehuber et al. 2010). The number of paralogous sHSP genes per genome can vary between 2 (average) in *Archea* and *Bacteria*, 3 (average) in fungi, 8 (average) in *Metazoa*, and up to 50 (maximum) in some plants (Kriehuber et al. 2010). Many vertebrates, including humans, contain ten paralogous sHSP genes, although some variation may occur, notably in the fish taxa. Among the vertebrate sHSPs, orthology to the human sHSPs can be easily established by phylogenetic analysis in most cases, whereas this is usually not possible across the other taxa.

Multiple sequence alignment (Fontaine et al. 2003; Franck et al. 2004; Kriehuber et al. 2010) reveals that the conserved αCD is flanked by regions of elevated sequence variability: the moderately conserved N-terminal region (NTR) and the highly variable C-terminal extension (CTE). Sometimes a short central region (CeR) with high sequence variability, positioned between the NTR and the αCD (possibly serving as a ‘hinge’ between both regions), is considered separately (Wang et al. 2000; Fontaine et al. 2003). This organization of the primary structure is summarized in Fig. S1. The αCD contains several β-strands forming two anti-parallel β-sheets (sandwich structure), whereas the flanking regions (NTR, CeR, CTE) are less ordered or even disordered (Carra et al. 2019; Webster et al. 2019; Boelens 2020). Whether or not such regions without a compact folding structure should be considered domains is debatable. Together, these domains and regions contribute to both the dynamic association of the monomers into high molecular mass complexes and to their holdase chaperone function.

The evolutionary history of sHSPs through all domains of life is characterized by certain unique features (Kriehuber et al. 2010): (1) According to the PFAM classification^3^, the overwhelming majority of sHSPs from all domains of life contain just one domain, the highly conserved αCD, contrary to other studied protein superfamilies which typically diverged by recruiting additional domains; (2) The evolution of sHSPs follows a ‘dimorphic pattern’: the αCD has a monophyletic origin reflecting the evolution of species, as opposed to the flanking regions which were historically remodeled several times in parallel but independent of the αCD. Thus, the αCD and the flanking regions have a fundamentally different evolutionary history. It is believed that this combination of the conserved αCD with the variable flanking regions (NTR, CeR, CTE) provides a unique and high degree of structural and functional speciation.

Four of these sHSP genes carry known mutations which have been associated with neuromuscular disease phenotypes in humans: HspB1 (Hsp27, Hsp25), HspB3, HspB5 (αB-crystallin), and HspB8 (Hsp22) (Datskevich et al. 2012; Benndorf et al. 2014; Vendredy et al. 2020), not counting polymorphisms or other sequence variants with unclear relation to disease. Mutant alleles of HspB1, HspB3 and HspB8 typically cause neuropathies with a spectrum of symptoms ranging from the clinical characteristics ‘Distal Hereditary Motor Neuropathy’ and ‘Charcot-Marie-Tooth Disease’ to some forms of ‘Amyotrophic Lateral Sclerosis’ and of myopathies (Table S1; Benndorf et al. 2014; Echaniz-Laguna et al. 2017; Adriaenssens et al. 2017; Katz et al. 2020; Chen et al. 2021), tentatively suggesting a functional overlap of these sHSPs. However, the pathogenicity of the HspB3 mutations has been recently disputed (Adriaenssens et al. 2017; Vendredy et al. 2020). Quite differently, mutations in HspB5 cause exclusively various forms of myopathies, including cardiomyopathies, and cataracts in the lens of the eye, the latter partially in association with the myopathies. The overwhelming majority among these disease-associated mutations are missense mutations, as opposed to a small number of identified frame shift, nonsense and elongation mutations (Benndorf et al. 2014; Vendredy et al. 2020). Disorders caused by the mutant sHSP alleles with their high penetrance belong to the group of Mendelian diseases (Quintana-Murci and Barreiro 2010), in spite of the fact that in most patients the symptoms develop only later in life (late onset diseases). Most of these missense mutations are associated with a dominant disease phenotype, or this association can be assumed, e.g., when the mutations occur sporadically (Table S1). Four missense mutations in HspB1 and HspB5 are associated with a recessive disease phenotype.

One approach for understanding the evolutionary forces that have shaped proteins is to measure the ratio (ω) of the non-synonymous (β) and synonymous (α) substitution rates (ω = β/α) at each codon position in a gene of interest (Kosakovsky Pond and Frost 2005). Non-synonymous substitutions change the amino acid being coded for at that site, which can directly affect protein structure and influence the fitness of an organism through negative (purifying) selective pressure. In contrast, synonymous substitutions do not change the amino acid being coded for, leaving the amino acid sequence unchanged. Synonymous substitutions are typically viewed as neutral and provide a baseline rate against which non-synonymous evolutionary rates can be calibrated. Therefore, the ratio ω of relative rates of non-synonymous and synonymous substitutions can provide information as to the type of selection that has acted upon a given set of protein-coding sequences, upon selected sequence segments (e.g. structural domains), or upon single codons (e.g. mutation sites). This ratio ω has become a standard measure of selective pressure in evolutionary biology (Frost et al. 2005; Arenas 2015). When there are more non-synonymous changes relative to synonymous changes, the ω ratio will be greater than 1, indicative of positive, diversifying selection. Conversely, if there are fewer non-synonymous changes relative to synonymous changes, the ω ratio will be less than 1, indicative of negative, purifying selection.

In this study we have estimated the ratio ω for each codon along the entire length of the sequences of human HspB1, HspB3, HspB5, and HspB8, using the aligned vertebrate sequences of the orthologs. We provide evidence for the exposure of the NTR, αCD, CeR, and CTE in these sequences to different degrees of purifying selection during vertebrate evolution. This disparate pattern of selective pressure within vertebrates fits well to the wider pattern of the dimorphic evolution that has shaped sHSPs across all domains of life, i.e., from the origin of the tree of life onward (Kriehuber et al. 2010). Thus, the same forces that shaped the sHSPs early in evolution are detectable also in the vertebrate evolution in recent 400-500 million years. Although similar, the ω profiles of the four studied human sHSPs exhibit differences indicating exposure of each sHSP to variable degrees of evolutionary pressure. For the disease-associated missense mutation sites, we find that all sites were historically exposed to detectable purifying selection, although to different degrees. Additionally, our data provide an explanation for the different incidence by which the affected sHSPs harbored disease-associated mutations.

## Methods

### Retrieval of vertebrate sHSP sequences from the databases

This study was restricted to the vertebrate sHSP sequences, specifically to the taxon *Gnathostomata*. *Gnathostomata* emerged more than 400 million years ago in evolution (Kuraku et al. 2016) and comprises all vertebrate species except those from the taxon *Agnatha*. *Gnathostomata* orthologs of human HspB1, HspB3, HspB5, and HspB8 were retrieved by a BLAST-P search at the NCBI website^4^, using the full-length human protein sequences as the query sequence. Additionally, several sequences were retrieved from the ENSEMBL website^5^, adding in total to sequences from at least 130 species for each studied sHSP, representing the major taxons of *Gnathostomata* (see below). Thereafter, the matching cDNA sequences were retrieved. Only full-length sequences of high quality were retained for analysis. When the translatable sHSP nucleotide sequences were embedded in larger DNA sequence blocks, the sHSP open reading frames were identified *in silico* by using the translation tool^6^.

### Inclusion criteria

Each retrieved candidate protein sequence was aligned to the human ortholog using the EMBOSS-Needle algorithm^7^, returning the degree of identity and similarity of each sequence to be tested with the human sequence. Sequences exhibiting ≥50% identity and ≥60% similarity were included in further analysis, with all other sequences being excluded. When aligned by the EMBOSS-Needle algorithm, a few candidate sequences exhibit a putative N-terminal extension, compared to the human orthologs and to the orthologs of nearly all other species. However, these putative N-terminal extensions were deduced *in silico* by algorithms based on DNA-sequencing data without any experimental support. Therefore, these putative N-terminal extensions can be assumed to result from an erroneous *in silico*-identification of the initiation codons and were consequently excluded from further processing. In these sequences, we used the canonical initiation codons based on the alignment with the respective human sHSPs.

Orthology of each candidate sequence with the human sHSPs was tested by constructing gene trees using alignment and phylogenetic inference (Kristensen et al. 2011) at the Clustal Omega web site^8^. Phylogenetic trees were constructed using all ten human sHSPs and the query sequence. Orthology was determined through analysis of patterns of sequence clustering within the reconstructed phylogenetic tree. Sequences were labeled as orthologs of human HspB1, HspB3, HspB5 or HspB8 if they grouped with these sHSPs, whereas sequences that did not group with these sHSPs were discarded.

Applying the criteria above resulted in the exclusion of several candidate sHSP-like sequences, notably from the fish taxa. The numbers of sHSP sequences that finally entered analyses for the detection of natural selection within the taxon *Gnathostomata* were 152 for HspB1, 130 for HspB3, 147 for HspB5, and 144 for HspB8 (Table S2).

### Species included in the sHSP sequence analysis

The designation of species and their taxonomic ranking was according to the NCBI web site^9^. This study includes sHSP sequences from *Gnathostomata* (*Vertebrata: Gnathostomata*) species representing the major vertebrate taxa: *Mammalia* (*Gnathostomata*: *Teleostomi*: *Euteleostomi*: *Sarcopterygii*: *Dipnotetrapodomorpha*: *Tetrapoda*: *Amniota*: *Mammalia*; includes primates, rodents, lagomorphs, ungulates, whales, bats, shrews, carnivorians, mammals of African origin, marsupials, armadillos, others), *Sauria* (*Amniota*: *Sauropsida*; *Sauria*; includes crocodiles, lizards, snakes, birds, turtles), *Amphibia* (*Tetrapoda*: *Amphibia*; includes frogs and toads), *Coelacanthimorpha* (*Sarcopterygii*: *Coelacanthimorpha*; includes just one species, the coelacanth), *Neopterygii* (*Euteleostomi*: *Actinopterygii*: *Neopterygii*, includes various taxa of ray-finned fish), and *Chondrichthyes* (*Gnathostomata*: *Chondrichthyes* or cartilaginous fish). The names of all species included in this study and their taxonomic ranking are given in Table S2.

### Determination of codon-specific evolutionary pressure along the entire length of the human sHSPs

The strength and direction of pervasive natural selection (i.e., purifying versus positive or diversifying) was assessed at the level of individual codon sites in each of the four sHSPs using the Fixed Effects Likelihood (FEL) method implemented in the HyPhy package (Pond et al. 2005) and the Datamonkey^10^ web application (Pond and Frost 2005). A dN/dS point estimate with confidence intervals was computed for each site using the FitMG94 model (Kosakovsky Pond et al. 2010). The confidence interval around the maximum likelihood point estimate of ω was derived by computing occurrences where the parameter estimate was not rejected in favor of the maximum likelihood estimate at a defined significance level (95% confidence of profile likelihood). A phylogenetic tree was constructed, using FastTree2.1 (Price et al. 2010) with the Generalized Time Reversible (GTR) model of nucleotide evolution. We estimated the ratio (ω) of non-synonymous (β) to synonymous (α) substitution rates for each codon along the entire multiple sequence alignment by maximum likelihood. These ω-values are a commonly used measure of selection, expected to be i) significantly (using the likelihood ratio test, LRT) less than one when purifying selection is predominant, ii) not significantly different from one for neutral evolution, and iii) significantly greater than one if positive selection is predominant. We report ω-value estimates for all codons of the human sHSP sequences and omit all codons in vertebrate sequences that have no homologous sites in the human sequences (Table S3). The estimated numeric ω-values also indicate the strength of the selective pressure for a given codon, e.g., with values of 0 or near 0 suggesting a greater degree of purifying selection. p-values indicating statistical significance were derived using the asymptotic chi-square test distribution for the LRT. Statistical significance (LRT p-value ≤ 0.1) for detecting purifying selection was observed for all codons with ω = 0 (no columns; cf. Fig. 1) and typically for most codons with 0 < ω < 0.5 (indicated by black columns). Statistical significance (LRT p-value ≤ 0.1) for detecting diversifying selection was observed for two codons in HspB3 with ω > 1.5 (indicated by blue columns). For codons with ω ≈ 1, no selective force was detected (neutral evolution; indicated by gray columns). Table S3 gives the numeric values of β, α, ω (dN/dS point estimates), and the dN/dS confidence intervals with the upper and lower bounds.

**Fig. 1.**
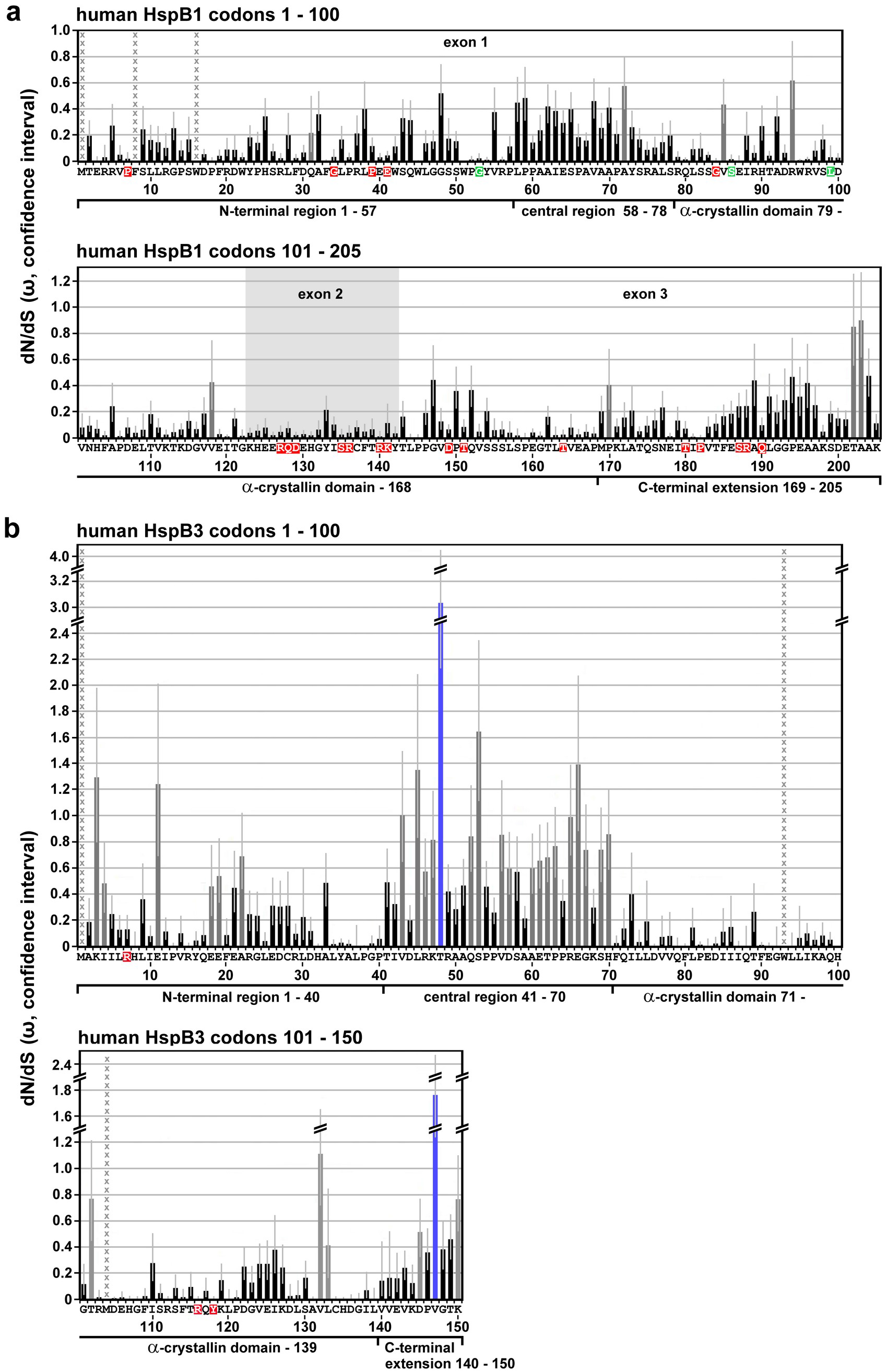
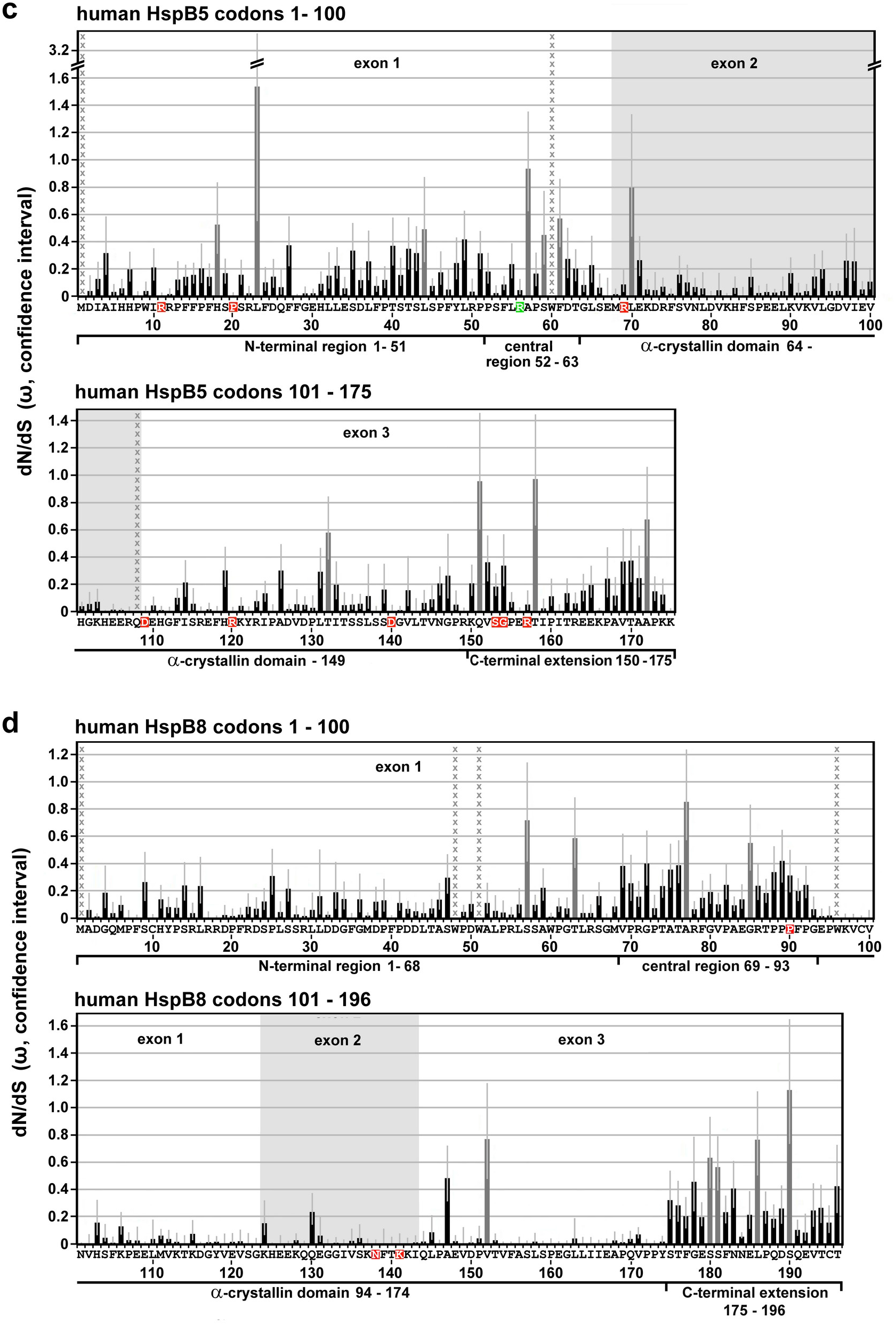
Selective pressures along the sequences of human HspB1 (a), HspB3 (b), HspB5 (c) and HspB8 (d) as returned by the FEL algorithm. The dN/dS point estimates were determined for each codon of the aligned *Gnathostomata* sHSP orthologs, omitting all codons without homologous amino acid residues in the human sequences. The plots show the principal ω-values with the respective confidence intervals (lower and upper bound; light gray error bars) along the primary sequences. No columns or black columns: ω < 1, indicating that these positions were historically exposed to purifying selection, using p ≤ 0.1 as statistical significance threshold; dark gray columns: ω ≈ 1 or near 1, with p > 0.1, indicating neutral evolution; blue columns: ω > 1, with p < 0.1, indicating positive selection. Positions with undefined ω-values are indicated by columns of gray x. Amino acid residues affected by disease-associated missense mutations are highlighted in color (red, green: dominant and recessive disease phenotypes, respectively). Sequence segments (partitions) with relatively low, intermediate or high ω-values in average, as they can be discerned, are indicated and were used to delineate the sequence partitions as used in this study (cf. Table S4). These partitions correspond approximately, though not perfectly, to the NTR, CeR, αCD, and CTE. The exons are indicated where applicable.

Because methionine is encoded solely by ATG, the ω-value for the strictly aligned (invariable) initiator methionines is not defined (β/α; β = 0, α = 0). Depending on the alignments, the ω-values for several internal methionine and tryptophan (encoded solely by TGG) residues, or occasionally for other amino acid residues within sequences may also be undefined (β/α; β = 0, α = 0) or approaching infinity (β/0; β/α with α → 0). These codons (marked in Table S3 and by columns of gray x in Fig. 1) which were included in the FEL analysis were excluded from the subsequent selection analyses using the FitMG94 method (see below), while other internal, variable methionine and tryptophan residues were included in both analyses.

### Sequence partitions

For comparison of the selective pressures acting historically on different parts of the sHSP sequences, the human sequences were separated into defined partitions. These partitions were primarily inferred from the profiles of the ω-values as returned by the FEL-method (cf. Fig. 1), thereby separating regions with relatively high, medium, or low average ω-values. The obtained partitions correspond approximately, though not perfectly, to the domains and regions as identified earlier by multiple alignment of the sHSPs (cf. Fig. S1): The NTR, αCD, CTE, and CeR can be discerned in most cases. Where this ω-plot-based discrimination was not possible (e.g., for some regions of HspB5), multiple alignment data were also used to demarcate the partitions as used in this study. Partitions corresponding to the exons of HspB1, HspB5, and HspB8 were separated according to the ENSEMBL web site^11^.

### Determination of the evolutionary pressure on full-length sequences and sequence partitions

In addition to the evaluation of selection at the level of individual codon sites, selection at the level of the coding regions of the sHSPs as a whole (full-length sequences) or at the level of the partitions (corresponding approximately to the NTR, CeR, αCD, CTE) was also evaluated. The goal of this partition-level selection analysis was to determine the selective pressure on defined regions relative to the other regions of that same sHSP. Additionally, the collected partition-level data permit the comparison of homologous partitions between the various sHSPs. Also, partitions corresponding to the exon structure of these sHSPs were compared to provide insight into the evolutionary history of the structure of these genes. Aggregate estimates of the omega values 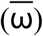 were calculated using the standard MG94xREV codon substitution model implemented in the FitMG94 algorithm^12^ (Kosakovsky Pond et al. 2010) and represent either the full-length sequence-wide inference or the partition-wide inference of the alignment-wide omega (ω) values (point estimates) with associated confidence intervals, an upper and lower bound. This approach provides a quantitative measure of the selective pressures historically applied to each partitioned region on average. The calculated alignmentwide aggregate 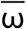-values are listed in Table S4 with upper and lower bounds, and are marked by indices corresponding either to the full-length sequences or to the various partitions, e.g. 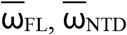, etc. To test for significantly different aggregate 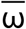 estimates, we performed a one degree of freedom likelihood ratio test where the null hypothesis enforced the same aggregate value on both genes or partitions, and the alternative hypothesis estimated them separately (FitMG94-Compare). Statistical relationships with significantly different full-length or partitioned sequences, as identified by this analysis, are indicated in the Hasse diagrams in Figs. 2 through 4 by arrows, pointing towards the sequences with the greater numeric aggregate 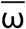 values, i.e., with less stringent purifying selection.

**Fig. 2.**
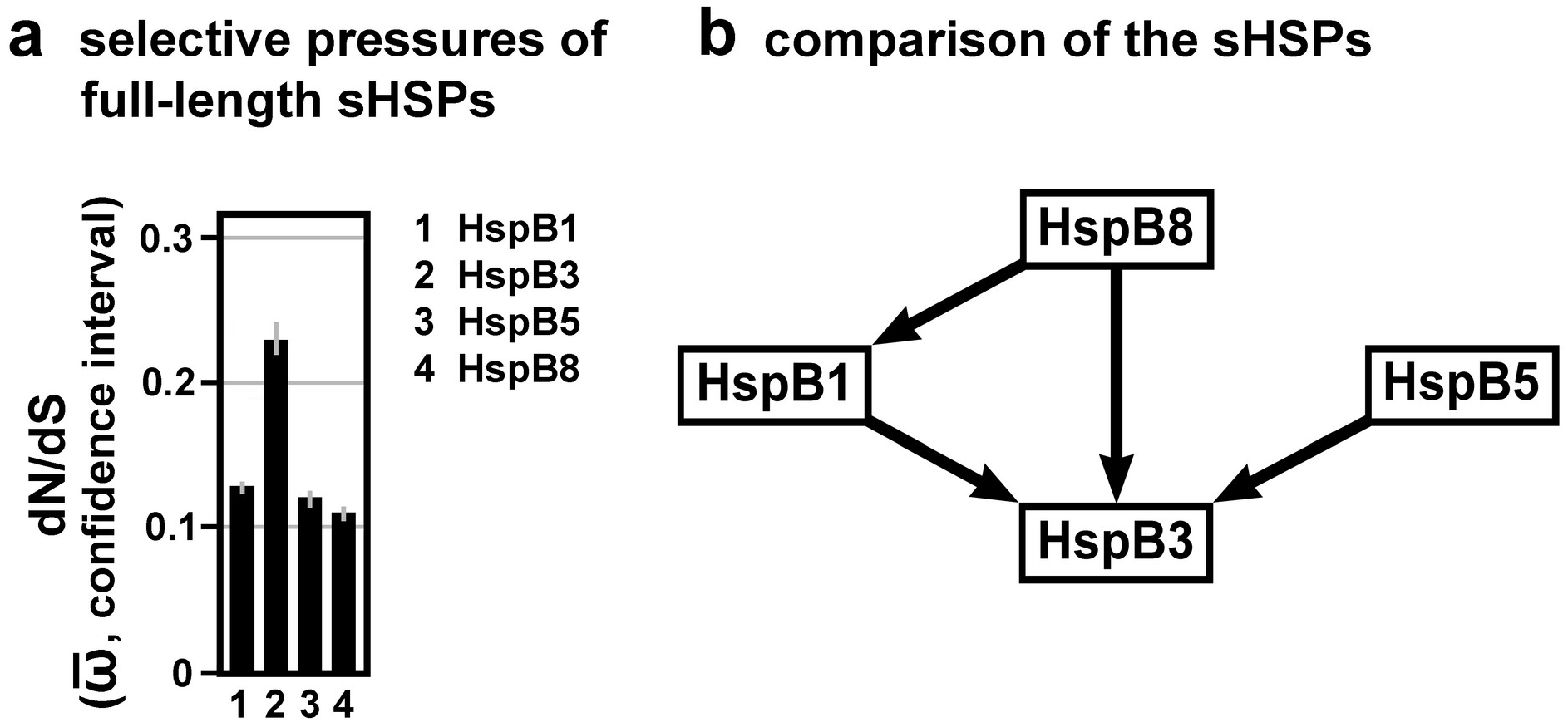
Selective pressures estimated for the full-length sHSP sequences. **a,** Plot of the dN/dS estimates (aggregate 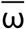-values with confidence intervals) of the full-length sHSPs, as returned by the FitMG94 algorithm. The data were taken from Table S4. **b,** Hasse diagram, as inferred by the FitMG94-Compare method, demonstrating the statistical relationship between the aggregate 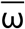-values of the four human sHSPs. The arrows point to the sHSPs with the higher 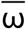-values, i.e., they were historically exposed to a less stringent purifying selection.

## Results

### Selective pressure profiles along the human sHSP sequences

The ω-values (dN/dS point estimates) for each codon of the *Gnathostomata* HspB1, HspB3, HspB5, and HspB8 alignments, as returned by the FEL model (Table S3), were plotted along the sequences of the human sHSPs, omitting all codons which have no homologous sites in the human sequences (Fig. 1). Overall, these results reveal that most of the codons were exposed to varying degrees of purifying selection over their evolutionary history (no columns, black columns). A minor fraction of codons in each sHSP do not reveal a detectable signal for purifying selective pressure, tentatively suggesting that they have evolved neutrally (gray columns). Obviously, there are differences between these sHSPs in terms of (1) the fraction of these neutrally evolving codons, (2) their distribution along the sHSPs sequences, and (3) the strength of the detected selective pressures (see below). In HspB1 the codons with neutral signature (8/205; 3.9%) are distributed relatively evenly along the sequence with just one cluster of two codons (T202, A203) found in the distal CTE (Fig. 1a). In HspB3 the fraction of neutrally evolving codons is highest among the studied sHSPs (28/150; 18.7%) with most of these codons clustered in the CeR (Fig. 1b). Interestingly, HspB3 is the only studied sHSP with codons that were historically exposed to a detectable positive diversifying selection (T48, V147), and not by chance these codons are found in regions with less stringent conservation or even neutral evolution: in the CeR and CTE. While in HspB5 the codons with neutral signature (11/175; 6.3%) are relatively evenly distributed over the entire sequence (Fig. 1c), in HspB8 these codons (9/196; 4.6%) are clustered, to a certain extent, in the CeR and CTE (Fig. 1d).

A closer inspection of FEL ω-plots (cf. Fig. 1) reveals alternating patterns with regions of lower and higher ω-values, indicating variable degrees of higher and lower purifying selection, respectively, acting over the evolutionary history of these sequence sections. Based on these profiles, the sequences of the human sHSPs were partitioned as specified in Fig. 1a through d (for HspB1, HspB3, HspB5, and HspB8, respectively) and in Table S4, in order to separate these sequence segments. These partitions correspond approximately to the domains and regions in the sHSPs as identified earlier by sequence alignments: NTR, αCD, CeR and CTE (cf. Fig. S1). Such differentiation of the ω-profile into regions with lower and higher ω-values is most pronounced in HspB3 (Fig. 1b) and HspB8 (Fig. 1d) and to a lesser extend in HspB1 (Fig. 1a). In HspB5 it is least pronounced (Fig. 1c) thus complicating the allocation of the partitions. In regions with unclear differentiation (i.e., low *vs*. high ω-values), information from the sequence alignment (Fontaine et al. 2003; cf. Fig. S1) was also used for the delimitation of the partitions. The partitions as defined in Fig. 1 and Table S4 were used to quantitatively determine the selective pressures (measured as aggregate 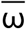) to which the respective regions of these sHSPs were historically exposed to (see below).

### Differences in the selective pressures between the sHSPs (full-length sequences)

Next, we attempted to quantitatively determine differences in the selective pressures acting over the evolutionary history of the *Gnathostomata* sHSPs, considering the full-length sequences. Based on the codon-specific ω-values (dN/dS point estimates) as shown in Table S3, aggregate 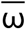-values for each full-length sHSP (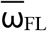; cf. Table S4) were inferred using the FitMG94 algorithm (Fig. 2a). Clearly, the obtained aggregate 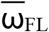-values between these sHSPs differ: HspB8 and HspB3 were historically exposed to the most stringent (lowest 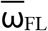) and most relaxed (highest 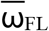) purifying selection, respectively, with HspB1 and HspB5 being placed in-between. Further statistical testing using the FitMG94-Compare algorithm confirms that several of these differences indeed exhibit a great statistical confidence, as represented by the Hasse diagrams (Fig. 2b). The arrows point to the sHSPs with higher aggregate 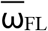-values: (1) HspB3 evolved under a less stringent purifying selection compared to the other studied sHSPs; and (2) HspB1 evolved under a less stringent level of purifying selection than HspB8. No differences were detected between HspB5 on the one hand and HspB8 and HspB1 on the other hand, applying the same statistical confidence criteria. Such differences in the overall selective pressures between some of the sHSPs, as demonstrated here, may be related to the frequency by which these sHSPs harbor disease-associated mutations (cf. Discussion).

### Differences in the selective pressures between the regions within the sHSPs

In the next step we sought to verify that different parts of the four studied sHSP sequences were indeed historically exposed to different degrees of selective pressures, as is suggested by Fig. 1. Again, we used the codon-specific ω-values (dN/dS point estimates; Table S3) and inferred, using the FitMG94 algorithm, the aggregate 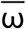-values for each sequence partition as defined in Fig. 1. As expected, the aggregate 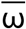-values for the various partitions exhibit differences within each sequence (Table S4). Plotting these partition-specific aggregate 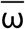-values results in profiles which are, to a certain extent, characteristic for the studied sHSPs (Fig. 3a). For example, the highest and least degrees of purifying selection, among all partitions, are seen in the αCD of HspB8 and CeR of HspB3, respectively, not counting the exon-specific partitions (see below).

**Fig. 3.**
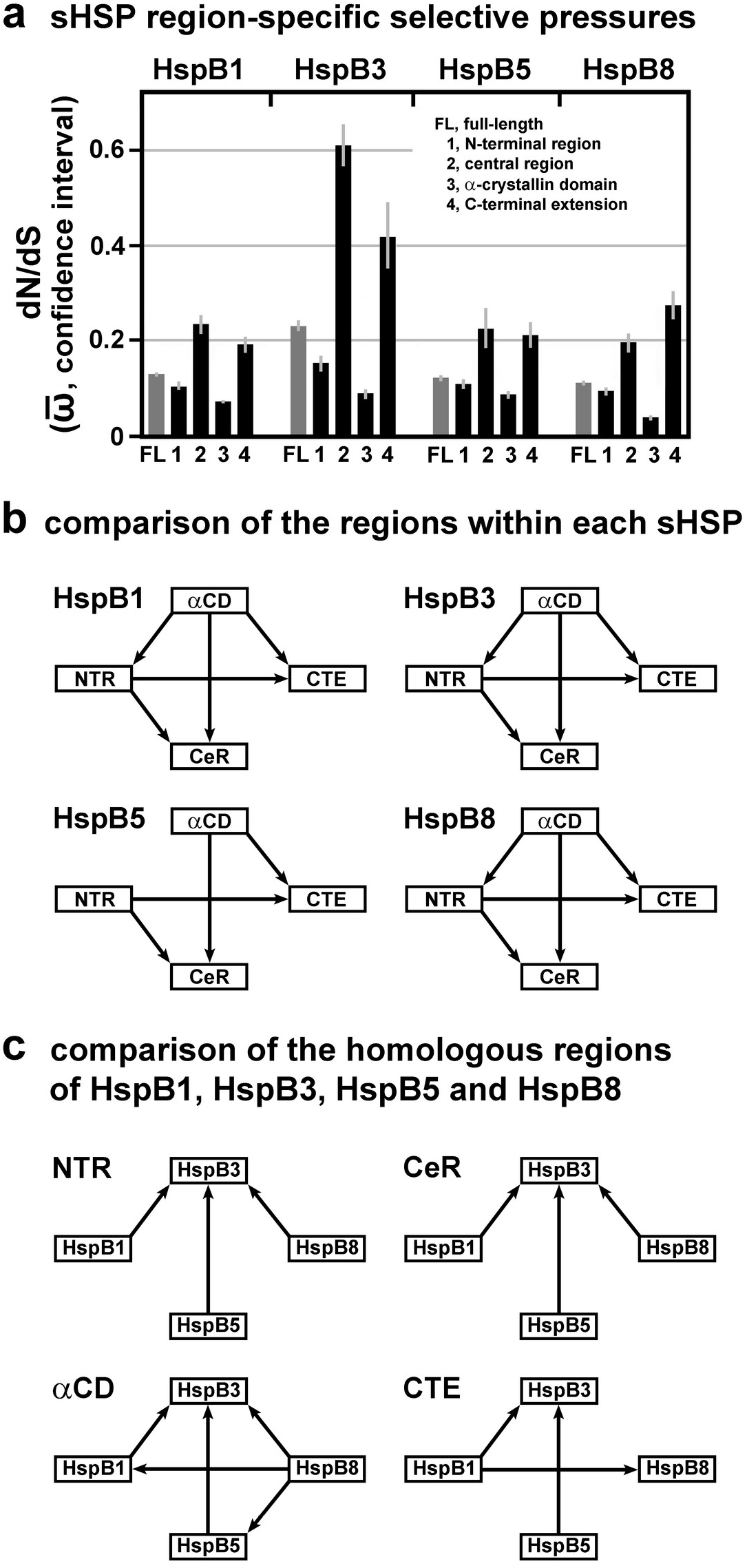
Selective pressures estimated for the various sHSP domains and regions. **a,** Plot of the dN/dS estimates (aggregate 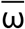-values with confidence intervals) of the region-specific partitions of the four studied sHSPs, as returned by the FitMG94 algorithm. The data were taken from Table S4. For comparison the aggregate 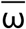-values of the full-length sequences are also included (gray columns). **b, c,** Hasse diagrams, as determined by the FitMG94-Compare method, demonstrating the statistical relationship between the aggregate 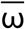-values of the regions within each sHSP sequence (b) or between the homologous regions of the four sHSPs (c). The arrows are used as in Fig. 2.

Further testing using the FitMG94-Compare algorithm reveals the high statistical confidence for most of the differences between the partitions within each sHSP sequence (Fig. 3b). The resulting Hasse diagrams demonstrate that the αCDs of HspB1, HspB3 and HspB8 were exposed to a more stringent purifying selection than all the other regions within the same sHSP sequences. For HspB5, however, no such statistical difference between the αCD and the NTR could be discerned, although the αCD was under a more stringent purifying selection than the CeR and CTE. In all four studied sHSPs, the NTR was under a more stringent purifying selection than the CeR and the CTE, whereas no differences were detected between the CeRs and the CTEs.

In summary, within each studied sHSP, sequence partitions can be identified which exhibit detectable and significant differences in the degrees of purifying selection they were historically exposed to.

### Differences in the selective pressures between the homologous regions of the four studied sHSPs

We used the collected aggregate 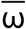-values (Table S4) also for the comparison among the homologous regions, as defined in Fig. 1 and Fig. S1, of the four studied sHSPs (Fig. 3a). Applying the FitMG94-Compare algorithm provided the Hasse diagrams shown in Fig. 3c, resulting in these relationships: The NTR, CeR, and αCD of HspB3 exhibit the least degrees of purifying selection when compared to the homologous partitions of the other sHSPs, while the CTE of HspB3 exhibits a lesser degree of purifying selection when compared to the homologous partitions of HspB1 and HspB5, but not of HspB8. The αCD of HspB8 was exposed to a stronger purifying selection than the αCDs of all other sHSPs. Quite differently, the CTE of HspB8 was exposed to a lesser degree of purifying selection than the CTE of HspB1. None of the partitions exhibits a significant difference between HspB1 and HspB5, a finding that is consistent with the fact that we could not detect a significant difference between the full-length sequences of both proteins (cf. Fig. 2b). Similarly, most of the partitions (NTR, CeR, CTE) do not exhibit a difference between HspB5 and HspB8, except the αCD which clearly exhibits a difference. This difference, however, does not have a sufficient weight to cause a difference at the level of the full-length sequences (cf. Fig. 2b). In summary, several significant differences in the degrees of purifying selection between the homologous domains and regions of the studied sHSPs were detected.

### Differences in the selective pressures between the exons

HspB1, HspB5 and HspB8 are encoded by genes consisting of three exons, whereas HspB3 is encoded by a one-exon gene (cf. Fig. 1). In HspB1, HspB5 and HspB8, exon 2 encodes the core part of the αCD, the most conserved region in these sHSPs (Fontaine et al. 2003; Franck et al. 2004). Applying the FitMG94 algorithm, we used the codon-specific ω-values (dN/dS point estimates; Table S3) and inferred the aggregate 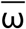-values for each exon-specific partition. As expected, plotting the exonspecific aggregate 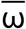-values for HspB1, HspB5 and HspB8 confirms that exons 2 were historically exposed to the strongest purifying selection, compared to exons 1 and 3, within each sHSP, with HspB5 exhibiting the least pronounced effect (Fig. 4a; cf. Table S4). When the FitMG94-Compare algorithm was applied, this finding was confirmed with a high statistical confidence (Fig. 4b). This analysis reveals also that exons 1 of HspB1 and HspB8 were under a more stringent purifying selection than exons 3 of the same sHSPs. In HspB5, no such difference between exons 1 and 3 can be demonstrated. When the homologous exons among the three sHSPs were compared, the FitMG94-Compare algorithm did not detect such differences which would hold out against this level of statistical confidence, with one exception: Exon 2 of HspB5 was historically exposed to a less stringent selective pressure than exons 2 of HspB1 and HspB8 (Fig. 4c). This relation is noteworthy and may have consequences for the frequency by which exons 2 in these sHSPs harbor disease-associated mutations (cf. Discussion). In summary, the various exons of HspB1, HspB5, and HspB8 were historically exposed to differential degrees of purifying selection. This relationship is most pronounced when exons 2 are compared with exons 1 and 3 within the same sHSP on the one hand and among one another on the other hand.

**Fig. 4.**
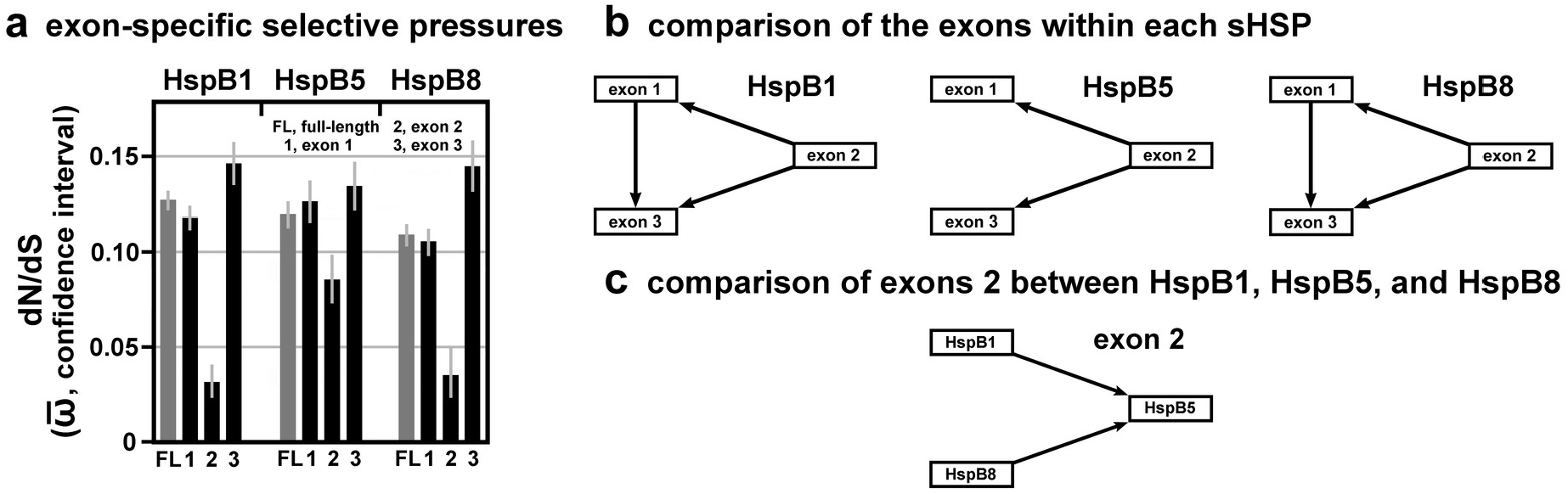
Selective pressures estimated for the exons of HspB1, HspB5, and HspB8. **a,** Plot of the dN/dS estimates (aggregate 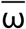-values with confidence intervals) of the various exon-specific partitions, as returned by the FitMG94 algorithm. The data were taken from Table S4. For comparison, the aggregate 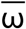-values of the full-length sequences are also included (gray columns). **b, c,** Hasse diagrams, as determined by the FitMG94-Compare method, demonstrating the statistical relationship between the aggregate 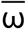-values of the exons within each sHSP sequence (b) or between exons 2 of all three sHSPs (c). The arrows are used as in Fig. 2.

### Selective pressure on the missense mutation sites

Based on the used set of *Gnathostomata* orthologs of HspB1, HspB3, HspB5 and HspB8, purifying selection was detected at all mutation sites harboring missense mutations (cf. Fig. 1, Table S3; with the relevant data summarized in Fig. 5). For most of these mutation sites, evidence for highly stringent purifying selection was found with dN/dS point estimates (ω-values) of zero or near zero (ω < 0.05). The range of the ω-values for the remaining mutation sites is < 0.35, indicating lesser stringent purifying selection. The four known mutation sites that harbored missense mutations which are associated with recessive disease phenotypes (G53, S86, L99 in HspB1; R56 in HspB5) all exhibit ω-values < 0.05, indicating highly stringent purifying selection.

**Fig. 5.**
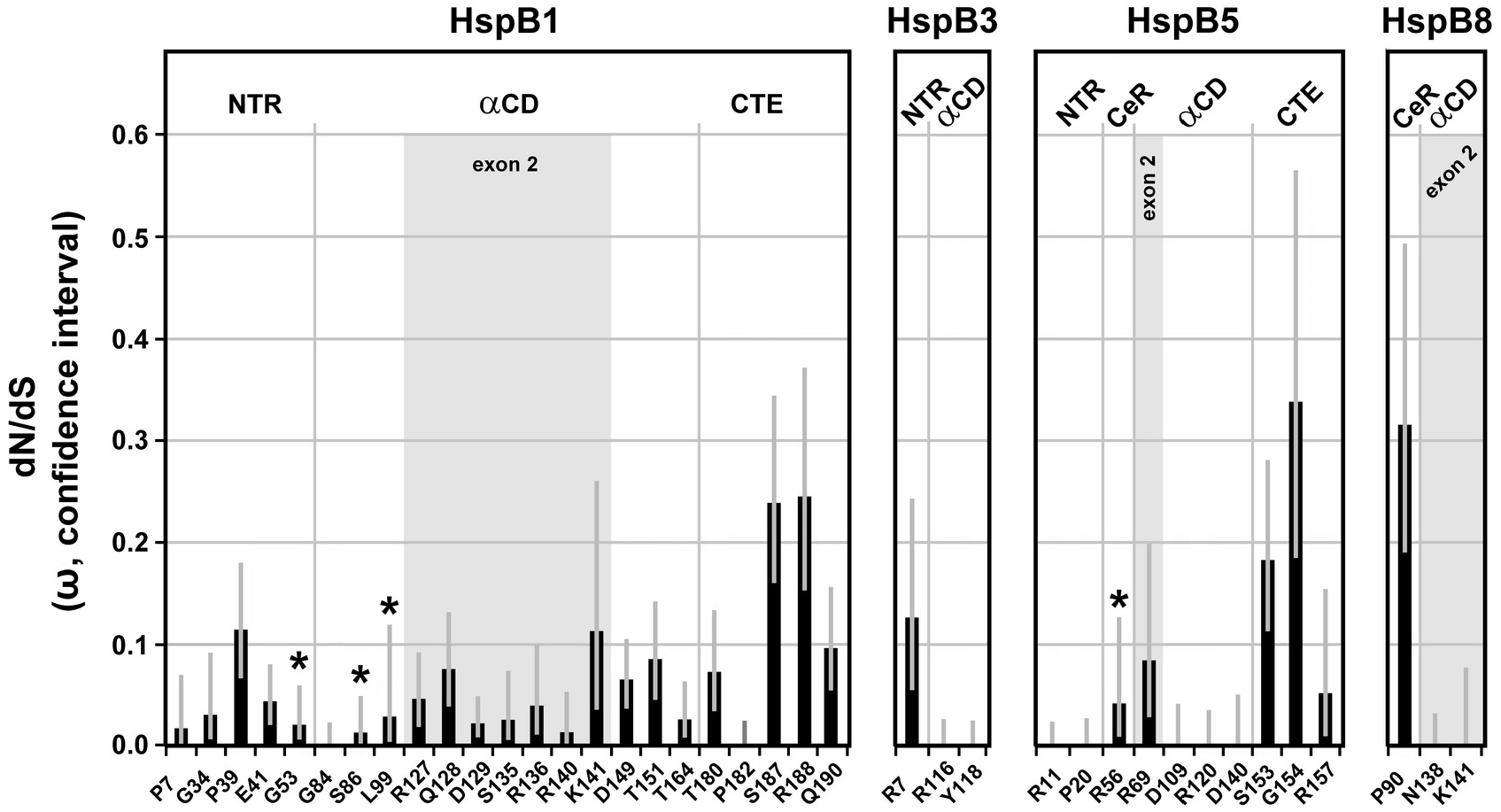
Summary of the selective pressures detected at the disease-associated missense mutation sites of the four studied sHSPs. The dN/dS point estimates (ω-values with confidence intervals) for each mutation site were taken from Table S3. The positions of the mutation sites within the various sequence partitions (domains, regions, exons) of the sHSPs are indicated (cf. Table S1, Fig. 1). The asterisks mark mutation sites whose mutations are associated with a recessive disease phenotype. All other mutation sites are affected by mutations that are associated with a dominant disease phenotype, or this association can be assumed.

Interestingly, there is a disparate distribution of the known disease-associated missense mutations among the exons in the various sHSPs: Exons 2 in HspB1 and HspB8 harbored a significant fraction of the known mutations in the respective sHSPs, whereas exon 2 in HspB5 harbored only a minor faction (just one mutation), among all mutations known to date (Fig. 5). The reason for this disparate distribution is not known, although it may be related to the relatively relaxed purifying selection seen in exon 2 of HspB5 (cf. Fig. 4; see Discussion).

## Discussion

In this study, we explore the evolutionary history of four of the ten human sHSPs: HspB1, HspB3, HspB5, and HspB8. These sHSPs are known disease factors as mutant alleles have been found associated with various forms of neuropathies, myopathies, and with cataracts in the lens of the eye. We found that the *Gnathostomata* sHSPs have been generally under strong purifying selection that is typical of highly conserved and functionally constrained genes. Interestingly, we observed varying degrees of purifying selection across the four sHSPs (highest: HspB8; lowest: HspB3; HspB1 and HspB5 being placed in between) (cf. Fig. 2). When the human sequences were partitioned into defined segments with relatively higher and lower selective pressures (corresponding approximately to the domains and regions of NTR, CeR, αCD, and CTE), our results demonstrate the highest degree of purifying selection in the αCD and less stringent purifying selection in the flanking regions, with the CeR and the CTE exhibiting the least degree of purifying selection. This finding is consistent with earlier comparative studies based on sequence alignments (Fontaine et al. 2003; Franck et al. 2004; Kriehuber et al. 2010). One finding of our study is that the obtained profiles of the strength of the selective pressures are characteristic attributes for the studied sHSPs.

Two codons in HspB3 (T48, V147) are distinguished such that they were historically under a clear positive, diversifying selection (cf. Fig. 1b). The reason for this observation is not known.

Concerning the exon structure, exons 2 of HspB1, HspB5, and HspB8 were subject to the most stringent purifying selection, compared to the other exons. If compared to one another, exon 2 of HspB5 was subject to the least stringent purifying selection (cf. Fig. 4). Interestingly, the regions of stringent purifying selection do not necessarily end at the borders of exon 2 and can extend into the nearby regions of exons 1 and 3. This is best seen with HspB8 (cf. Fig. 1d).

With regard to the disease-associated mutations sites, all of them were found to be under stringent purifying selection (cf. Fig. 5). Two further observations are noteworthy: (1) All mutation sites in the αCD of HspB3 and HspB8, and three out of four mutations sites in the αCD of HspB5, exhibit only synonymous variation (ω = 0), as opposed to most mutation sites in the αCD of HspB1 that exhibit both, synonymous and non-synonymous variations; and (2) mutation sites positioned in the CTEs of HspB1 and HspB5 exhibit, by trend, weaker purifying selective pressure than mutation sites in the other partitions.

Our results provide a partial explanation for some of the unusual features of the sHSPs, including the dimorphic pattern of their evolution, the observed differences in the mutation rates between the sHSPs in humans, and the prevalence of genetically dominant disease phenotypes resulting from gain-of-function mutations. These features are discussed in detail below:

(1) *The dimorphic pattern of sHSP evolution*. As determined earlier by phylogenetic analysis (Kriehuber et al. 2010), the evolutionary history of the sHSPs with the dimorphic pattern exhibits a striking feature: While the isolated central part of these proteins, the αCD, evolved along the species tree across all domains of life from the beginning, the flanking regions (NTR + CeR, CTE) have trees different from the αCD being consistent with extensive remodeling and complex evolutionary dynamics independently of the αCD. The same analysis revealed also that all *Metazoa* (including *Gnathostomata*) sHSPs are derived from a single ancestral sequence, with the gene number increasing by gene duplication mostly after the separation of the different *Bilateria* taxa during the Cambrian species explosion. Interestingly, certain characteristic N-terminal sequences found otherwise scattered all over the αCD-based phylogenetic tree and which served to establish the evolutionary dimorphic pattern, were not detected in the *Metazoa* branch of this phylogenetic tree (Kriehuber et al. 2010). In this context, our collected data provide an argument for the extension of this dimorphic evolutionary pattern also to the branch of the *Gnathostomata* sHSPs. Differences in the selective pressures (measured as aggregate 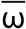-values) between the αCD with its most stringent purifying selection on the one hand and the flanking regions (NTR + CeR, CTE) on the other hand with their more relaxed purifying selection imply that this dimorphic evolutionary pattern continued also in the taxon *Gnathostomata* (cf. Fig. 3b).

A closer inspection reveals also certain differences in the selective pressures between the different partitions of the flanking regions in all four studied sHSPs. For example, the purifying selective pressures in the NTR was historically stronger than in the CeR and CTE of all four studied sHSPs (cf. Fig. 3b). These differential selective pressures likely played a role in shaping the vertebrate sHSPs, even if the specifics are not understood at this time. The importance of these flanking regions for both the structure and function of the mammalian sHSPs has been demonstrated (Selig et al. 2020).

(2) *Differences in the mutation rates between the sHSPs in humans*. As shown in Table S1 and in the references therein, sHSPs harbor disease-associated mutations with different incidences. This is especially evident if HspB1, HspB3, and HspB8 are compared, all of them causing, when mutated, a similar spectrum of peripheral motor neuropathies. HspB1 harbored by far most of the mutations as compared to HspB3 and HspB8: In a systematic study of a cohort of 510 unrelated index patients with peripheral neuropathies, 32 patients were found to have mutations in sHSPs (32/510; 6.3%) (Echaniz-Laguna et al. 2017). Among them, 28 patients carried mutations in HspB1, four patients in HspB8, and none in HspB3. Thus, in the patient population of this study, HspB1 is approximately seven times more frequently affected than HspB8, while HspB3 apparently plays only a marginal role, if any. In another study of 163 patients belonging to 108 families with distal hereditary motor neuropathies, a genetic diagnosis was achieved in 37/108 families (34.2%) or in 78/163 of all patients (47.8%) (Frasquet et al. 2021). Among them, the most frequent cause of disease was mutations in HspB1, besides mutations in several other genes. In that study, no mutations were found in HspB8 or HspB3. Similarly, in other systematic studies with smaller patient cohorts, a number of mutations in HspB1 were found, but none in the other sHSPs (Capponi et al. 2011; Luigetti et al. 2016).

This disparate incidence may be directly related to the different degrees of purifying selection these sHSPs have been historically subjected to (measured as aggregate 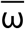-values for the full-length sequences 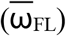). HspB3 and HspB8 were exposed to the most relaxed and most stringent purifying selection, respectively, with HspB1 being placed in-between (cf. Table S4, Fig 2). Because of the historically most stringent purifying selection, it is plausible that mutations in HspB8 would lead more frequently to impaired fitness, thus eliminating the disease alleles over time from the human population. Conversely, mutations in HspB3 may have fewer consequences for the fitness, thus allowing the accumulation of mutated alleles in the population. If this were the case, there should exist a relatively large number of HspB3 allelic variants in the healthy human population. HspB1, positioned in-between, exhibits a certain degree of ‘plasticity’, allowing it to harbor mutations at an increased rate compared to HspB8, although this comes with a cost for the health. However, since a significant proportion of mutant HspB1-associated disorders have a late onset in life (references in Table S1), these mutations are likely to come with relatively little fitness costs in the human population in evolutionary terms. In this scenario, most mutant HspB1 alleles should be under weak purifying selection in the human population, if any, thus permitting their accumulation.

These considerations cannot simply be extended to HspB5 as the disease phenotypes associated with its mutations are quite different. However, in terms of both the estimated overall selective pressure for full-length HspB5 (cf. Fig 2) and the number of the harbored missense mutations (cf. Fig. 5), HspB5 may share some similarity with HspB1. Yet, another feature is noteworthy: Exon 2 of HspB5 harbored only a minor fraction (just one) of the known mutations, quite different from the exons 2 of HspB1 and HspB8 that harbored a major fraction of the known mutations (cf. Fig. 5). This may be related to the fact that exon 2 of HspB5 is distinguished by the relatively relaxed purifying selection it was historically subjected to, compared to HspB1 and HspB8 (cf. Fig. 4a, c). Following the above logic, exon 2 of HspB5 may have a relatively high tendency to accumulate a number of allelic variants in the healthy human population without major fitness costs in evolutionary terms.

*(3) The prevailing of genetically dominant disease phenotypes resulting from* gain-of-function *mutations*. The fact that most missense mutations in sHSPs are associated with dominant (or semi-dominant) disease phenotypes is contrary to the expectation, as mutagenesis studies in many organisms revealed that ~90% of the wild-type alleles are dominant over the mutant alleles, with dominant mutant phenotypes being the exception in the wider picture (Wilkie 1994). Thus, in addition to the peculiar dimorphic evolutionary history, sHSPs are also distinguished by their atypical genotype-phenotype relation.

Heritable diseases in general are believed to persist in the human population because of a balance between mutations, genetic drift, and natural selection, the latter eventually eliminating the mutation from the population by purifying selection (Blekhman et al. 2008). For alleles causing highly penetrant Mendelian disorders, purifying selection can be quite strong, unless the disease outbreak has a late onset as is the case with most mutant sHSP-associated disorders (see references in Table S1). Thus, in the human population the majority of the mutant sHSP alleles can be expected to be under reduced purifying selection, as the late onset is unlikely to impose major fitness costs in evolutionary terms. This is, however, not in conflict with the fact that all sHSP mutation sites themselves were exposed to a stringent purifying selection, together with the entire genes, during the *Gnathostomata* evolution, as we show here. Indeed, it was observed that Mendelian disease genes in general are under a more widespread and stronger purifying selection if the disease-causing alleles are dominant, as compared both to recessive and more complex disease genes (Blekhman et al. 2008). While this interrelation is highly interesting, the mechanisms behind it still await their elucidation.

Disease-causing mutated alleles of the four studied sHSPs with their high phenotypic penetrance and segregation of the disease traits in a Mendelian fashion qualify them as Mendelian disease genes. All four sHSP genes harboring such mutations were historically subject to strong purifying selection. These facts and observations offer an explanation for both the high proportion of dominant mutations among neuromuscular disease-associated sHSP mutations, and also for the relatively high proportion of sHSP mutations (6.3% and 10.4%; Echaniz-Laguna et al. 2017; Frasquet et al. 2021; respectively) among all mutations in patients with inherited peripheral neuropathies.

Historically the phenomenon of genetic dominance has been fiercely debated (summarized in Benndorf et al. 2014). One aspect is that the existence of multiple paralogous genes, such as the 10 sHSP genes in humans, seems to be related to their receptivity to harboring dominant-negative mutations. The evolutionary retention of such paralogs in polyploid species, or species with polyploid ancestry that includes *Homo sapiens*, was explained by gene dosage effects and dominant-negative effects involving supramolecular structures (Comai 2005; Veitia 2010; Veitia and Birchler 2010). Such oligomeric complexes are indeed a hallmark of all studied sHSPs (Mymrikov et al. 2011; Boelens 2020; Muranova et al. 2020).

Mutations associated with dominant phenotypes fall into two categories: loss-of-function type (i.e., haploinsufficiency) and gain-of-function type (Wilkie 1994), although mixed types might also occur. For the dominant sHSP mutations, a loss-of-function situation can be tentatively dismissed. Arguments come from HspB1-, HspB5-, and HspB8-knock-out rodents that do not reproduce overt disease phenotypes as are seen for the dominant missense mutations (Brady et al. 2001; Huang et al. 2007; Bouhy et al. 2018), despite the fact that some impaired cell biological pathways, e.g. the autophagosome formation, could be identified (Haidar et al. 2019). Additional biochemical evidence supports this notion: Mutations in sHSPs do not necessarily result in the loss of key functions such as chaperoning and anti-apoptotic activities (Krishnan et al. 2008; Almeida-Souza et al. 2010), even if some modification, e.g. of the chaperoning specificity, may occur (Weeks et al. 2018). These findings tentatively suggest that loss-of-function mechanisms may not be the primary mechanism responsible for the manifestation of the disorders.

Regarding the dominant gain-of-function mutations, there are only a handful principal mechanisms that seem to accommodate most situations (Wilkie 1994): (1) ‘classic’ dominant negative effects (e.g. through protein-protein interactions); (2) formation of toxic gene products (e.g. of amyloids); (3) impact on cytoskeletal systems resulting in altered cell architecture; and (4) increased enzymatic activities. Existing experimental, genetic and clinical evidence suggests that most if not all studied dominant disease-associated sHSP mutant alleles fit this pattern. Table S5 summarizes some of the molecular and cellular consequences of the expression of dominant mutant sHSP alleles, all supporting gain-of-function mechanisms. They result in abnormal protein-protein interactions and in abnormal quaternary structures (implying dominant negative effects, summarized e.g. for a number of HspB1 mutants by Muranova et al. (2020)), in the deposition of protein aggregates (implying the formation of highly toxic amyloid precursors and altered cell architecture), and in increased downstream enzymatic activities. This situation should have direct consequences for any future therapeutic interventions which should aim to reduce these toxic effects of the mutated proteins, e.g. by reducing the expression of the mutant proteins, by ‘detoxification’ of amyloid precursors of the aggregates, or by inhibition of the abnormally increased downstream enzymatic activities. The latter point was experimentally addressed by d’Ydewalle et al. (2011) who found that expression of mutant HspB1 decreased the abundance of acetylated α-tubulin in mice that was paralleled by axonal transport deficits. A pharmacological inhibition of the histone deacetylase 6 restored the α-tubulin acetylation together with the axonal transport, and thus rescued the disease phenotype. Also, pharmacological targeting sHSPs and their abnormal interactions with other proteins resulting from mutations is within reach of future approaches (Boelens 2020).

Taken together, our data shed light onto some unusual features of the evolution of four of the vertebrate sHSPs (HspB1, HspB3, HspB5, HspB8), and how their evolutionary history has shaped these proteins. We provide evidence that the vertebrate sHSPs exhibit a dimorphic evolutionary pattern, with different degrees of purifying selection acting historically upon the αCD on the one hand and on the flanking regions on the other hand. When compared to one another, the four studied sHSPs have been exposed to different degrees of purifying selection. These features have implications for the role of these sHSPs in human health and disease and may explain the different incidences by which the various sHSPs harbored disease-associated mutations in the human population and also the unusual genotype-phenotype relation in patients with neuromuscular diseases resulting from missense mutations in these sHSPs. Experimental findings and theoretical considerations support the prevalence of toxic gain-of-function mechanisms for the manifestation of the mutant sHSP-associated diseases in most individuals. We propose that any future therapeutic approaches should directly tackle these toxic properties of mutant sHSPs, or of toxic downstream consequences of their expression. Examples include the interference in abnormal protein-protein interactions involving mutant sHSPs, the inhibition of the formation of toxic amyloids, the protection of the cytoskeletal architecture, or the inhibition of abnormally increased downstream enzymatic activities.

Our study has demonstrated how the analysis of the evolution of vertebrate genes, which cause inheritable diseases if mutated, has the potency to lead to future medical strategies. In so doing, we provide support for Theodosius G. Dobzhansky’s famous realization that ‘nothing in biology makes sense except in the light of evolution’ (Dobzhansky 1973). The extension of this paradigm to inheritable human diseases may help to advance the emerging field of ‘Evolutionary Medicine’.

## Footnotes

3 PF00011 at http://pfam.xfam.org/family/PF00011#tabview=tab0

4 https://www.ncbi.nlm.nih.gov/

5 https://useast.ensembl.org/index.html?redirect=no

6 https://web.expasy.org/translate/

7 https://www.ebi.ac.uk/Tools/psa/emboss_needle/

8 https://www.ebi.ac.uk/Tools/msa/clustalo/

9 https://www.ncbi.nlm.nih.gov/taxonomy/

10 https://www.datamonkey.org

11 https://useast.ensembl.org/index.html?redirect=no

12 https://github.com/veg/hyphy-analyses/tree/master/FitMG94

